# Spatial Multimodal Analysis of Transcriptomes and Metabolomes in Tissues

**DOI:** 10.1101/2023.01.26.525195

**Authors:** Marco Vicari, Reza Mirzazadeh, Anna Nilsson, Reza Shariatgorji, Patrik Bjärterot, Ludvig Larsson, Hower Lee, Mats Nilsson, Julia Foyer, Markus Ekvall, Paulo Czarnewski, Xiaoqun Zhang, Per Svenningsson, Per E. Andrén, Joakim Lundeberg

## Abstract

We present a spatial omics approach that merges and expands the capabilities of independently performed *in situ* assays on a single tissue section. Our spatial multimodal analysis combines histology, mass spectrometry imaging, and spatial transcriptomics to facilitate precise measurements of mRNA transcripts and low-molecular weight metabolites across tissue regions. We demonstrate the potential of our method using murine and human brain samples in the context of dopamine and Parkinson’s disease.

## Main

Spatially resolved transcriptomics (SRT) allows the measurement of genome-wide mRNA expression and also provides positional information about the mRNA in a tissue section. Although key aspects of SRT technologies can vary, each technique will ultimately yield a gene-expression count table with tissue coordinates^1–3^. Mass spectrometry imaging (MSI) enables label-free, spatially-resolved measurement of the abundance of biomolecules directly from fresh frozen tissue sections^4,5^. In matrix-assisted laser desorption/ionization (MALDI)- MSI, a matrix is applied to the surface of tissue sections mounted onto a glass slide. Focusing a pulsed laser beam onto the tissue section then generates ionic species from the molecules present in the sample surface, enabling the collection of mass-to-charge (m/z) spectra at defined raster positions from across the tissue section. Although the SRT and MSI technologies are becoming more popular in spatial biology, they are currently applied as separate methodologies due to experimental constraints such as non-charged, barcoded (in SRT) versus conductive (in MALDI-MSI) microscopy slides, along with the risk of RNA degradation during the harsh MSI process. In the present paper, we demonstrate the possibility of combining SRT and MALDI-MSI in a single tissue section with retained specificity and sensitivity of both modalities, by introducing a spatial multimodal analysis (SMA) protocol.

SMA workflow comprises four steps: i) sectioning non-embedded snap-frozen samples onto non-charged, barcoded gene expression arrays, ii) mass spectrometry imaging by MALDI, iii) hematoxylin and eosin (H&E) staining and bright field microscopy, and iv) SRT (**Fig. 1A**). To test the feasibility of our method, we assessed whether RNA transcripts remain intact after tissue exposure to MALDI-MSI by using a slide coated with polydT probes. We mounted coronal sections of a mouse brain and sprayed it with four different MALDI matrices: 1) 9-aminoacridine (9-AA) for detection of metabolites in negative ionization mode, 2) 2,5-dihydroxybenzoic acid (DHB) for detection of metabolites in positive ionization mode, 3) norharmane for detection of various lipids, and 4) 4-(anthracen-9-yl)-2-fluoro-1-ethylpyridin-1-ium iodide (FMP-10), which charge-tags molecules with phenolic hydroxyls and/or primary amines, including neurotransmitters^6^. We imaged the sections using Fourier-transform ion cyclotron resonance (FTICR)-MS, and collected spectra for approximately three hours at RT. Fluorescence microscopy imaging of the cDNA footprint generated during the RNA quality check assay showed that the captured transcripts correlated well with tissue morphology, indicating that mRNA, surprisingly, is still present after MALDI-MSI in all of the investigated matrices (**Suppl. Fig. 1A-C**). The presence and integrity of mRNA post-MALDI were confirmed using targeted *in situ* sequencing^7^ on coronal mouse sections with FMP-10 on conductive MALDI slides (**Suppl. Fig. 1D**).

**Figure 1.**
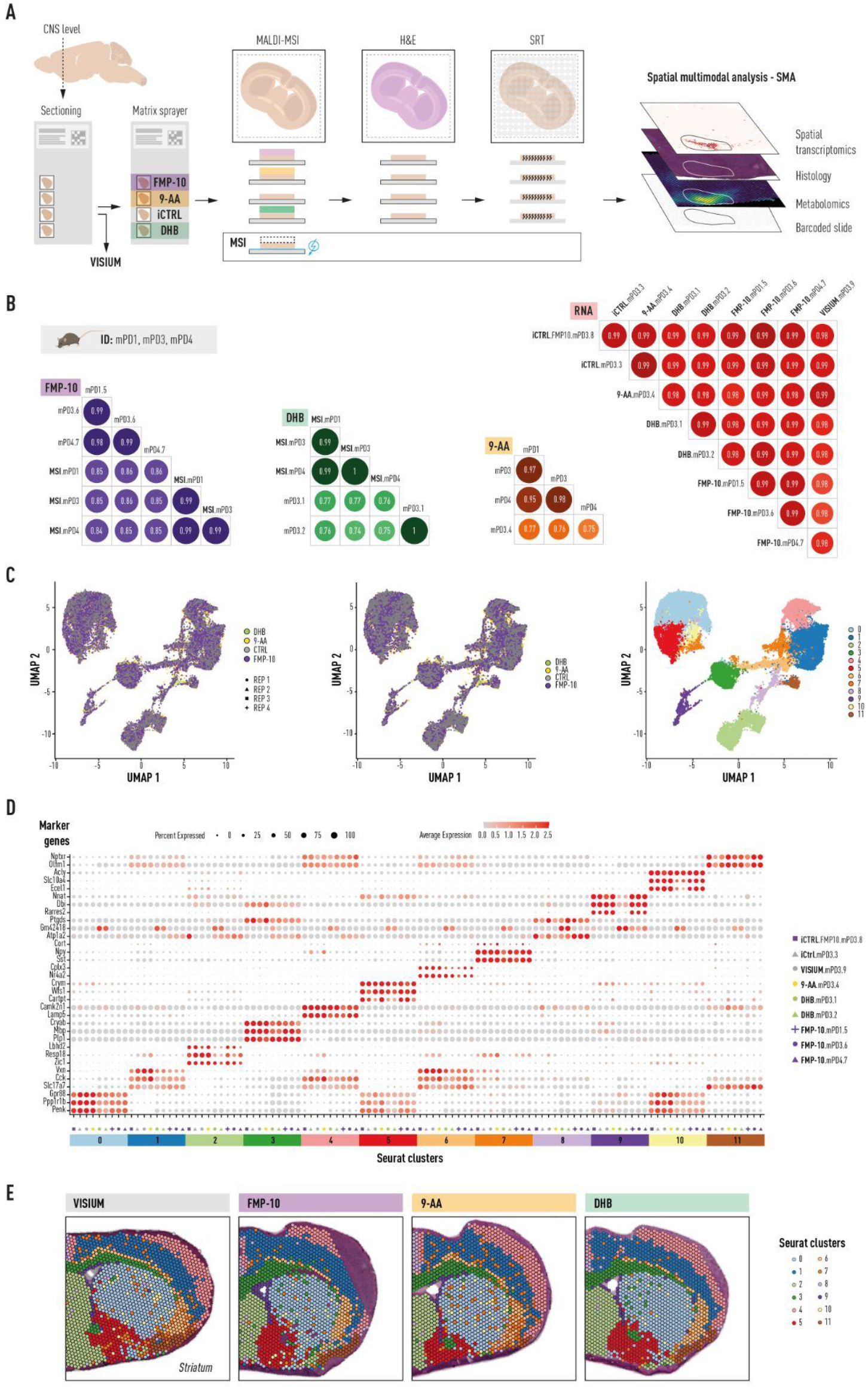
A multimodal spatial omics approach to investigate metabolites, morphology and gene expression analysis. **A)** The SMA workflow: non-embedded, snap-frozen samples are sectioned and thaw-mounted onto non-charged, barcoded Visium gene expression arrays. Tissue sections are then sprayed with MALDI matrices and MSI is performed. This is followed by H&E staining and imaging with bright field microscopy. Finally, sections are processed for SRT. **B)** Pairwise gene-to-gene and molecule-to-molecule correlations across biological and technical replicates. Samples are named with short identifiers that reflect the technical conditions under which the sample was analyzed: *MSI*: standalone MALDI-MSI, *Visium*: standalone Visium, identifiers that begin with the name of a MALDI matrix (FMP-10, 9-AA or DHB) in the gene-to-gene correlation plot (far right) indicate that the SMA protocol was used and which matrix was applied on the section. Serial numbers at the end of the identifiers represent which mouse (mPD1, mPD3 or mPD4) was used, and the serial number of the tissue section. **C)** Uniform manifold approximation and projection (UMAP) of SMA ST spots colored by sections (left), MALDI matrices (middle) and clusters (right). **D)** Top three marker genes with highest log2 fold change of each spatial clusters across technical conditions and biological replicates. **E)** Spatial plot of mouse brain tissue sections (striatal level, 0.49 mm from bregma) which illustrates clusters of transcripts for samples sprayed with three different MALDI matrices (FMP-10, 9-AA, DHB) and one sample processed with the standalone Visium protocol.

Next, we investigated the reproducibility of SMA. For this purpose, we repeated the experiments using barcoded Visium oligonucleotide slides, which enabled the quantification of individual captured transcripts by sequencing. We used seven consecutive coronal mouse brain sections from three different mice, imaged with three different MALDI matrices (9-AA, DHB, and FMP-10). We compared gene expression and MALDI-MSI data from matching tissue sections that had been analyzed either with the standalone Visium or MSI (ITO conductive slide), respectively. SMA data analysis demonstrated that the gene expression and small molecule profiles correlated well with the reference data produced using the two corresponding methods individually; the average number of unique genes per spot ranged between 2034 and 4001, and all of the calculated correlations were above 0.74 and p-values were lower than 0.05 (**Fig. 1B, Suppl. Figs. 2-4**). Joint analysis and visualization of transcriptomics data showed that the gene expression integrates well across experimental conditions, with similar expression of marker genes and high conservation of the spatial clusters (**Figs. 1C-E**). These results provide strong evidence that SMA can be used with several matrices for simultaneous gene expression and biomolecule profiling.

To further demonstrate the applicability of SMA, we used it on a mouse model of Parkinson’s disease (PD). PD is the most common neurodegenerative disorder among the human population after Alzheimer’s disease^8,9^. It is characterized by the loss of dopaminergic neurons within the substantia nigra pars compacta (SNc), containing neurons that project to the dorsal putamen of the striatum^10^. In the present study, we aimed to capture both gene expression and corresponding neurotransmitters from two brain regions (SNc and striatum) of three unilateral 6-hydroxydopamine (6-OHDA)-lesioned mice^11^ (**Fig. 2A, Fig.2Bi, Fig.2Bvi, Suppl. Fig. 5**). As expected, SMA predominantly detected dopamine in the intact striatum and SNc, but not in the lesioned contralateral hemisphere (**Fig. 2Bii, Fig. 2Bvii**). The multimodal data produced through SMA was also leveraged to identify the expression of genes associated with dopamine expression. Unsurprisingly, the key dopaminergic pathway genes *Th, Slc6a3, Slc18a2* and *Ddc* were found to be correlated with dopamine expression in the SNc^12^. Likewise, in the striatal sections, the dopamine levels were positively correlated with the expression of genes like *Pcp4* and *Tac1*, and negatively correlated with genes like *Penk* and *Cartpt*, suggesting dysregulation of medium spiny neurons (MSNs) (**Fig. 2Biv, Fig. 2Bix, Fig.2Di-ii**), while corroborating previous findings^13–15^. To substantiate our results, we sought to perform cell type deconvolution^16^ using scRNA-seq data from a mouse brain atlas^17^. Strikingly, we found a lower proportion of midbrain dopaminergic neurons MBDOP2 in the lesioned SNc and ventral tegmental area (VTA). A similar phenotype was observed in the lesioned dorsal striatum for MSN1 neurons, a subtype of medium spiny neurons (**Fig. 2Bv, Fig. 2Bx**). Furthermore, we were able to specify the localization of multiple neurotransmitters and metabolites, such as taurine, 3-methoxytyramine (3-MT), 3,4-dihydroxy-phenylacetaldehyde (DOPAL), 3,4-dihydroxyphenylacetic acid (DOPAC), norepinephrine, serotonin, histidine, tocopherol and GABA **(Suppl. Fig. 5**). The results demonstrated a similar spatial distribution of molecules as was previously reported in rat models using MALDI-MSI alone^6^.

**Figure 2.**
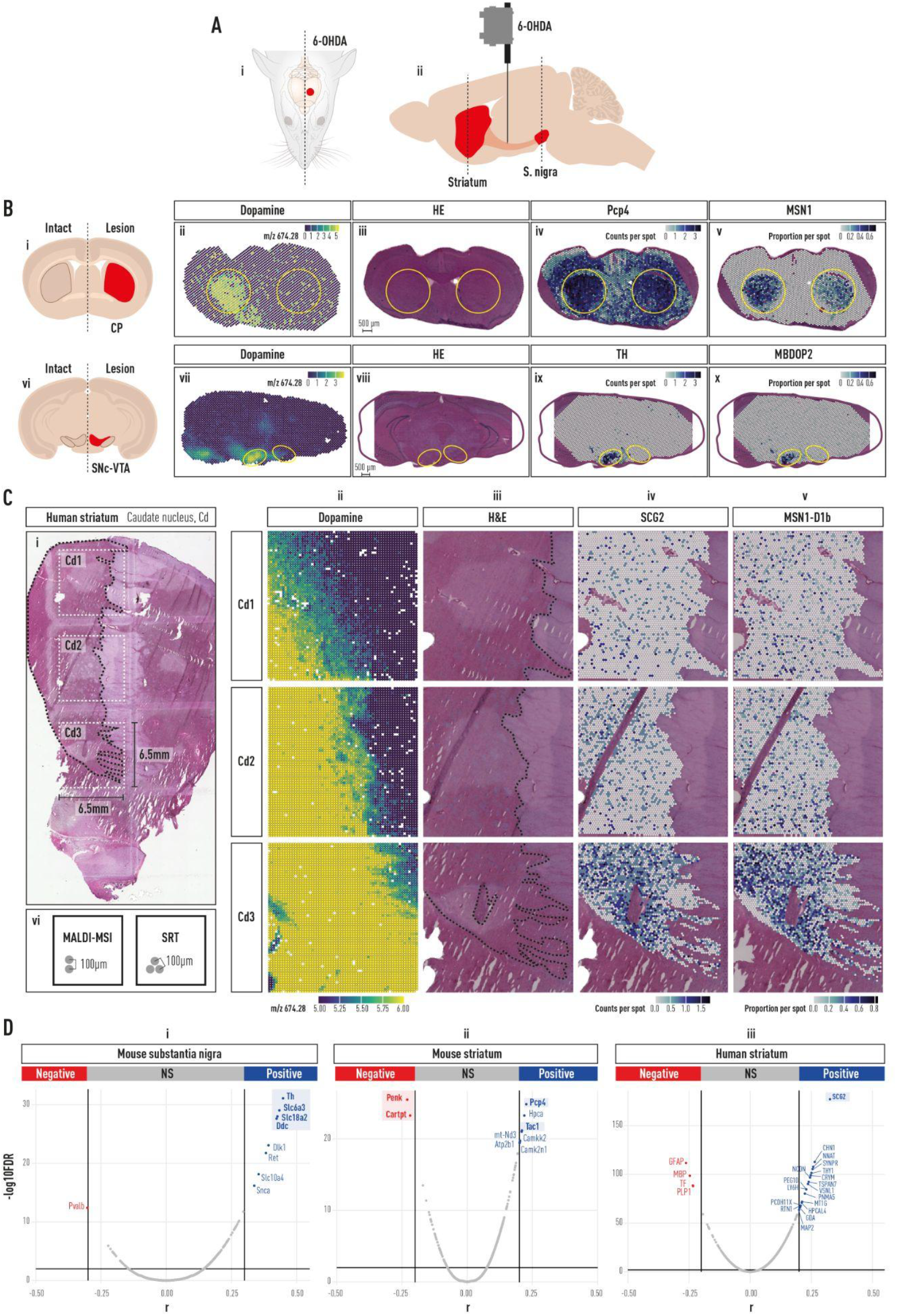
Spatial multimodal analysis of a Parkinson’s disease mouse model and a human post-mortem brain affected by Parkinson’s disease. **A)** Cartoon showing the injection of 6-OHDA only in one hemisphere in the median forebrain bundle (MFB). Dashed lines indicate the depth (0.49 and -3.39 mm, distance from bregma) for the substantia nigra and striatum, respectively. **B)** Representative sections from the substantia nigra and striatum of the mouse PD model. From left to right: cartoon showing the dopamine-depleted regions (panels i and vi), dopamine expression (panels ii and vii), H&E staining (panels iii and viii), spatial gene expression of the gene with highest correlation to dopamine (panels iv and ix), proportions of MSN1 (panel v) and MBDOP2 (panel x). **C)** Human post-mortem striatum sample. From left to right images are presented in the same order as in B. The demarcated area indicates the caudate nucleus of the striatum. See figure 2 for gene counts statistics. **D)** From left to right: dopamine-to-gene correlations in the mouse substantia nigra, striatum, and human caudate nucleus. SNc - substantia nigra pars compacta, VTA - ventral tegmental area (VTA), CP - caudoputamen, Cd - Caudate nucleus.

To demonstrate the relevance of the presented multimodal approach in human specimens, we applied it to a frozen human PD post-mortem striatal brain sample and measured neurotransmitters and gene expression over a 2.4 × 0.5 cm tissue section (**Fig. 2Ci**). To address and overcome RNA degradation in post-mortem material, we applied a recent protocol specifically developed for gene expression measurements from fresh frozen tissue samples with low/moderate RNA quality^18^. The spatial MSI distribution of DA and 3-MT (**Fig. 2Cii, Suppl. Figs. 6**) confirmed our previous findings^6^ in that these neurotransmitters were observed at higher levels in the medial division of the ventral caudate nucleus. Multimodal correlation analysis identified *SCG2* as the transcript which was most correlated with dopamine abundance (**Fig. 2Civ, Fig.2Diii**). We performed cell-type deconvolution analysis using publicly available snRNAseq data from the caudate nucleus of human post-mortem samples^19^, which contains six clusters of MSN neurons (MSN.D1a-c, MSN.D2a-c). Interestingly, similar to our observation in the PD model, MSN.D1b neurons were enriched in the dopamine-expressing region, with a spatial pattern similar to *SCG2* (**Fig. 2Cv**).

To summarize, we present a method that enables the simultaneous spatial profiling of small molecules and gene expression within a tissue section. The impact of this technology could be extended to other disciplines including oncology, where it could provide new insights into the dynamic crosstalk that regulates the tumor microenvironment^20^ and drives the response to treatment^21^. Gene expression can also be leveraged to infer genomic integrity^22^, which is important to matching tumor clones with drug efficacy. The presented approach provides a further level of multimodality when studying small molecules in a tissue context.

## Methods

### Animal Experiments

A total of four adult male C57Bl/6J mice, 8 weeks old (Charles River, Sulzfeld, Germany), were housed under controlled temperature and humidity (20 °C, 53% humidity) with a 12 h light/12 h dark cycles. The mice had access to standard food pellets and water ad libitum. All of the animal work was performed in agreement with the European Council Directive (86/609/EE) and approved by the local Animal Ethics Committee (Stockholms Norra Djurförsöksetiska Nämnd, no. 3218-2022).

During the experiments, one mouse served as the control while three mice were anesthetized with isoflurane (Apoteket, Stockholm, Sweden), pre-treated with 25 mg/kg desipramine intraperitoneally (i.p.) (Sigma–Aldrich, St. Louis, MO, USA) and 5 mg/kg pargyline i.p. (Sigma–Aldrich), placed in a stereotaxic frame, and injected over 2 min, with 3 μg of 6-OHDA in 0.01% ascorbate (Sigma–Aldrich) into the median forebrain bundle (MFB) of the right hemisphere over two minutes. The coordinates for injection were anterior-posterior (AP) −1.1 mm, medial-lateral (ML) −1.1 mm, and dorsal-ventral (DV) −4.8 mm relative to bregma and the dural surface^23^. During the post-operative phase, the analgesic buprenorphine (Temgesic 0.1 mg/kg) was subcutaneously (s.c.) was administered for two days following surgery. Two weeks after unilateral 6-OHDA administration, the lesion was validated by administering the mice with 1 mg/kg apomorphine i.p. (Sigma–Aldrich) and assessing rotational behavior in the mice. After the mice were sacrificed, the brains were removed, collected, and stored at -80 ºC for further use.

Efforts were taken to minimize the number of animals used and their suffering. Animals were euthanized by decapitation and brains were rapidly removed, snap-frozen in dry-ice cooled isopentane for 3 seconds, and stored at −80 °C to minimize post-mortem degradation.

### Human post-mortem sample

The human post-mortem sample was from the caudate-putamen level of the brain (coronal sections) of a man who died at 94 years of age. The post-mortem interval until the brain was frozen was 9.25 h. The neuropathological diagnosis was Parkinson’s disease in Braak stage 3. The case was obtained from the Harvard Brain Tissue Resource Center at the McLean Hospital (Belmont, MA, USA). Analyses were approved by the local ethical committee (Karolinska Institutet, Stockholm, Sweden, no. 2014/1366-31). All experiments were performed in compliance with all relevant ethical regulations.

### Tissue Processing and Sample Preparation

Coronal mouse brain tissue sections, 12 μm thick, were cut at −20 °C using a CM1900 UV cryostat-microtome (Leica Microsystems, Wetzlar, Germany) and subsequently thaw-mounted onto Visium glass slides (10x Genomics, Pleasanton, CA, USA) (for SMA and Visium analysis) or conductive indium tin oxide-coated glass slides (Bruker Daltonics, Bremen, Germany) (for MSI analysis). Sections were collected at the striatal level (distance from bregma, 0.49 mm^23^) and at the substantia nigra level (distance from bregma -3.39mm). The human striatal PD sample was sectioned at 12 µm thickness and the caudate region was placed over the four printed areas on the Visium slide (**Suppl. Fig. 6**). The prepared slides were stored at −80 °C. Sections were desiccated at room temperature (RT) for 15 min prior to scanning on a flatbed scanner (Epson Perfection V500, Nagano, Japan), with the exception of tissues coated with FMP-10, which were scanned after matrix application. For neurotransmitter analysis, on-tissue chemical derivatization was performed with the FMP-10 reactive matrix according to a previously described protocol^6^. Briefly, a freshly prepared solution of FMP-10 (4.4 mM) in 70% acetonitrile was sprayed onto mouse brain tissue sections and the human tissue sample over 20 passes at 90 °C using a robotic sprayer (TM-Sprayer; HTX Technologies, Chapel Hill, NC, USA) with a flow rate of 80 μL/min, spray head velocity of 1100 mm/ min, 2.0 mm track spacing, and 6 psi nitrogen pressure. Tissue sections from the control mouse and from one lesioned mouse were also coated with either 9-AA (5 mg/mL dissolved in 80% methanol; for analysis in negative ionization mode) or 3,5-dihydroxybenzoic acid (DHB, 35 mg/mL dissolved in 50% acetonitrile and 0.2% TFA; for analysis in positive ionization mode). 9-AA was applied using the TM-sprayer (75 °C, six passes, solvent flow rate of 70 μL/min, spray head velocity of 1100 mm/min, and track spacing of 2.0 mm), while DHB was applied using the same settings, except for a nozzle temperature of 95°C. Norharmane (7.5 mg/ml in 80% MeOH) was sprayed in 16 passes using the TM-sprayer with the following parameters: temperature 60°C, flow rate of 70 μL/min, spray head velocity of 1200 mm/ min, 2.0 mm track spacing, and 6 psi nitrogen pressure. Tissue sections mounted on the same glass slide but coated with different matrices were masked using a glass cover slip.

### MALDI-MSI

Tissue sections were imaged at 100 µm lateral resolution using a MALDI-FTICR (Solarix XR 7T-2Ω, Bruker Daltonics, Germany) instrument equipped with a Smartbeam II 2 kHz Nd:YAG laser. Laser power was optimized at the start of each analysis. Spotted red phosphorus was used for external calibration of the methods. Spectra were collected by compiling the signals from 100 laser shots per pixel. Samples coated with FMP-10 and DHB were analyzed in positive ionization mode. The quadrupole isolation *m/z* ratio (Q1) was set at *m/z* 379 (FMP-10) or *m/z* 150 (DHB), and data were collected for samples coated with FMP-10 and DHB over the *m/z* 150−1050 range and *m/z* 129-1000 ranges, respectively. For the FMP-10 analysis *m/z* 555.2231 was used as the lock mass and the matrix peak at *m/z* 273.0394 used as lock mass for internal *m/z* calibration of the data acquired from the DHB coated sample. Samples coated with 9-AA were analyzed in negative ionization mode over the *m/z* 107.5-1000 range with a Q1 mass of *m/z* 120 and *m/z* 193.0771 was used as the lock mass. Immediately after MSI analysis, the tissue containing glass slides with tissue samples were washed twice for 30s in pre-chilled methanol and then stored at -80 °C until Visium gene expression/tissue optimization processing.

### MALDI-MSI and SMA Metabolomics Data Analysis

The SCiLS Lab API (Bruker Daltonics) was used to create ion images that would be used in downstream analyses. To ensure similar *m/z* lists among samples with different derivatization matrices, a reference peaklist was used to calibrate all of the samples with the same derivatization matrix; thus, there was only one list of *m/z* values per matrix. To measure the MSI performance on Visium and ITO glass slides, Pearson correlations of mean spectra from consecutive sections analyzed on ITO or Visium were calculated using the SCilS Lab API (Bruker Daltonics) and the python programming language. UMAPs were performed in the R programming language with a similar script that was used for spatial transcriptomics UMAP, but modified to accomodate MSI data.

### Visium Spatial Gene Expression and Tissue Optimization

FF samples were cryo-sectioned at 10 µm thickness, mounted onto Visium glass slides and stored at -80°C before processing. Spatial gene expression libraries were generated following 10x Genomics Visium Gene Expression and Tissue Optimization protocols according to the manufacturer’s recommendations (*Visium Spatial Gene Expression Reagent Kits - Tissue Optimization User Guide*, Document Number CG000238 Rev E, 10x Genomics, (February 2022); *Visium Spatial Gene Expression Reagent Kits - User Guide*, Document Number CG000239 Rev F, 10x Genomics, (January 2022), *Methanol Fixation, H&E Staining & Imaging for Visium Spatial Protocols*, Document number CG000160 Rev C, 10x Genomics). Libraries were sequenced using a NextSeq2000 sequencing system (Illumina, San Diego, CA, USA). The length of read 1 was 28 bp, while the length of read 2 was 150 bp long.

### SMA

Sections of the Visium Gene Expression or Tissue Optimization glass slides (10x Genomics) were desiccated at RT for 15 min before the reactive matrices were applied.

Matrix application and MSI were performed as already described in the section *MALDI-MSI*. After MSI, the slides were briefly immersed in pre-chilled methanol (a total of three times), followed by storage at -80°C until Visium gene expression/tissue optimization was performed. Visium Spatial Gene Expression and Tissue Optimization slides, with the exception of the human post-mortem sample, were processed according to the corresponding latest versions of the 10x Genomics protocols (*Visium Spatial Gene Expression Reagent Kits - Tissue Optimization User Guide*, Document Number CG000238 Rev E, 10x Genomics, (February 2022); *Visium Spatial Gene Expression Reagent Kits - User Guide*, Document Number CG000239 Rev F, 10x Genomics, (January 2022), *Methanol Fixation, H&E Staining & Imaging for Visium Spatial Protocols*, Document number CG000160 Rev C, 10x Genomics), without any modification. Libraries were sequenced using the Nextseq2000 sequencing system (Illumina, San Diego, CA, USA). The length of read 1 was 28 bp, while the length of read 2 was 150 bp long.

The human post-mortem sample was processed according to the RRST protocol^18^. The slide was taken out of the -80°C freezer and placed on a thermocycler, pre-heated to 37°C, for 1 minute, followed by immediate fixation in 4% methanol-free formaldehyde (ThermoFisher, Waltham, MA, USA; Catalog number: 28906) solution for 10 minutes, RT. After fixation, the slide was washed twice in 1xPBS, heated up at 37°C for 20 minutes on a thermocycler, cooled down to RT, stained with Hematoxylin and alcoholic Eosin and imaged with a light microscope. After imaging, the slides were washed with MQ water, air-dried and placed inside the plastic Visium cassette. Sections were treated with 0.1N HCl for 1 minute at room temperature, and washed once in PBS. The Visium Spatial Gene Expression for FFPE reagent kit (10x Genomics, Pleasanton, CA, USA) was used in the subsequent steps. The de-crosslinking step was skipped, which means that probe pre-hybridization step (15 minutes RT) was followed by probe hybridization (occurring overnight) according to the 10X Visium Spatial Gene Expression reagent kit for FFPE protocol and library preparation (User Guide, CG000407 Rev C). Final libraries were sequenced using a Nextseq2000 (Illumina, San Diego, CA, USA). The length of read 1 was 28 base pairs, while the length of read 2 was 50 base pairs.

### Visium data processing

Sequenced libraries were processed using Space Ranger software (version 1.2.1 for standard Visium data and version 1.3.1 for RRST data, 10x Genomics). Reads were aligned to the pre-built human or mouse reference genome, including a GTF file, a fasta file and a STAR index, provided by 10x Genomics (GRCh38 for human data or mm10 for mouse data, version 32, Ensembl 98).

### Visium, RRST and SMA transcriptomics Data Analysis

The spatial transcriptomics data obtained with either standard Visium, RRST or SMA were processed and analyzed using R (v4.1.3), the single-cell genomics toolkit Seurat and the spatial transcriptomics toolkit STUtility. The H&E images were manually annotated based on tissue morphology and dopamine expression using the interactive application Loupe Browser (10x Genomics). Mouse striatum and substantia nigra hemispheres were categorized into two groups, i.e., “intact” for the left hemisphere and “lesioned” for the right. The filtered count matrices obtained from spaceranger were used in subsequent analysis upon application of additional filters. In particular, spots below sectioning or mounting artifacts were annotated using Loupe Browser and removed using the “SubsetSTData” function in STUtility; spots that included more than 38% mitochondrial genes or less than 50 unique genes were removed using the same STUtility function; hemoglobin-coding, riboprotein-coding and Malat1 genes were removed from the dataset as well. Pearson correlation coefficients and p-values were calculated using the continuous= “cor” argument in ggpairs function and the corrplot function of the *GGally* or *corrplot* R packages, respectively. After filtering out spots and genes as previously described, the data were normalized and subjected to a basic analytical workflow using functions from the Seurat R package. The SCTransform function was used for normalization and variance stabilization, and was followed by dimensionality reduction via PCA (RunPCA). Data were integrated with the RunHarmony function from the *harmony* R package using group.by.vars = “Sample.ID” (which indicates the sample of origin), assay.use = “SCT” and reduction = “pca” as parameters. A shared nearest neighbor (SNN) graph was constructed based on the first 30 principal components (FindNeighbors). Finally, a uniform manifold approximation and projection (UMAP) embedding was computed based on the first 30 principal components (RunUMAP); this was followed by graph-based clustering (FindClusters). Marker genes for each identified cluster were calculated using the function FindAllMarkers with default parameters, while a non-parametric Wilcoxon rank-sum test and the Bonferroni correction were used for p-value adjustment. Only genes with log2 fold change higher than 0.25 and adjusted p-value lower than 0.01 were considered differentially expressed. MSI and SRT were manually aligned using the interactive Shiny application (available in STUtility) through the function ManualAlignImages. The alignment procedure requires images of the tissue sections to be aligned - one image for the RNA data and one image for the MSI - as input. However, since we only had access to H&E images for the RNA data, we modified the alignment procedure to align the MSI data points directly to the corresponding H&E image. Prior to alignment, the MSI data first underwent PCA in order to identify and remove data points that were located outside of the tissue sections. Once the RNA data had been aligned to the MSI data, we identified pairs of nearest neighbors across the two datasets using the kNN function from the *dbscan* R package, with k set to 5. To remove parts of the tissue sections that did not overlap in the two datasets, we filtered out pairs with a distance higher than 35 pixels, a threshold that was empirically determined from the histogram of neighbor distances. Next, only the closest neighbor in the RNA data was kept for each MSI data point, thus generating a list MSI-RNA data point pairs. As MSI yielded a lower density of data points than Visium, we decided to select neighbors for MSI data points rather than the other way around. In cases where the multiple MSI data points shared the same nearest neighbor, this neighbor was reused to ensure a one to one mapping. Once the MSI-RNA data point pairs had been identified, we used these pairs to subset the raw data and produce new Seurat objects with the two aligned data modalities stored in separate assays. Identification of the genes that were most correlated with dopamine levels was performed by calculating pairwise Pearson correlation coefficients between dopamine and all of the genes retained in the dataset. P-values were adjusted using Benjamini and Hochberg correction method, and only genes with a false discovery rate lower than 0.01 were retained. Cell type proportions were inferred using stereoscope, a probabilistic method designed to deconvolve spatial data using single cell data. These analyses were run in accordance with the developer’s recommendations (https://github.com/almaan/stereoscope), using the 5000 most highly variable genes, based on the FindVariableFeatures function from the Seurat R package. The spatial transcriptomics data of the mouse model was deconvolved using scRNAseq data from the mouse brain atlas (http://mousebrain.org/adolescent/). Cells occuring more than once in the single cell count matrix were removed from the dataset. The 39 annotated taxa were used as a basis for deconvolution, along with the 4 annotated clusters of dopaminergic neurons and the 6 annotated clusters of medium spiny neurons present in the dataset. 50000 epochs were used for both single cell and spatial transcriptomics data, and the batch sizes were set to 2048. Single cell data was subsetted to a maximum of 1000 cells per cell type, using the --sc_upper_bound option. The spatial transcriptomics data of the human sample was deconvolved using snRNAseq data from GSE178265 repository. This dataset was filtered to retain only caudate nucleus samples, high quality nuclei (number of genes between 1000 and 10000, number of UMIs lower than 50000 mitochondrial genes content lower than 7%) and a subsample of 4000 cells per donor. The data were then normalized and subjected to a basic single cell analysis workflow. The functions SCTransform and RunPCA from the Seurat R package were used for normalization and variance stabilization, and dimensionality reduction, respectively. Data were integrated with the RunHarmony function from the harmony R package using group.by.vars = “orig.ident” (which indicates the sample of origin), assay.use = “SCT” and reduction = “pca” as parameters. The functions FindNeighbors, RunUMAP and FindClusters from the Seurat R package were used to construct an SNN graph, compute a UMAP embedding and perform graph-based clustering. Using a resolution of 0.6, a total number of 23 clusters were detected and used for cell type deconvolution. 75000 epochs were used for both single cell and spatial transcriptomics data, and the batch sizes were set to 100. Single cell data was subsetted to a maximum of 250 and a minimum of 25 cells per cell type, using the -sub and -slb options respectively. After deconvolution, the cell type proportion values were overlaid on the tissue section images by using the FeatureOverlay function in the STUtility package.

### Targeted ISS sample pretreatment

Post MALDI imaging, the mouse coronal section was fixed with 4% Formaldehyde for 5 min, followed by permeabilization with 0.1M HCl for 5min. The section is then dehydrated in an ethanol series of 70% and 100% for 2 min each.

### Targeted ISS Protocol

PLP Hybridization was performed overnight at 37C with 10 nM final concentration of phosphorylated padlock probes in PLP hybridization buffer (2X SSC, 10% Formamide). Sections were then washed 2x with PLP hybridization buffer and 2x with PBS. Ligation was performed for 2h at 37C with 1x T4 Rnl2 Reaction buffer (NEB B0239SVIAL), 1 U/µl T4 Rnl2 (NEB #M0239), 1 U/µl RNase Inhibitor (BLIRT #RT35) and RCA Primer at final concentration of 50nM. The sections were washed 2x with PBS before proceeding with rolling circle amplification at 30C, overnight with 0.5 U/μl Φ29 polymerase (Monserate Biotech #4002) in reaction mixture of 1X Φ29 buffer (50mM Tris-HCl, 10mM MgCl_2_, 10mM (NH_4_)_2_SO_4_), 5% Glycerol, 0.25 mM dNTPs (BLIRT #RP65), 0.2 μg/μl BSA.

Fluorescent probe detection was performed by hybridization of readout detection probes (100 nM) and DAPI (Biotium #S36936) in hybridization buffer (2X SSC, 20% Formamide) for 45 mins at RT. Sections were washed with PBS and mounted with SlowFade Gold Antifade Mountant (Thermo Fisher Scientific #S36936).

### Imaging of Targeted ISS

All images were obtained with a Leica DMi8 epifluorescence microscope equipped with an external LED light source (Lumencor® SPECTRA X light engine), automatic multi-slide stage (LMT200-HS), sCMOS camera (Leica DFC9000 GTC), and objectives (HC PL APO 10X/0.45; HC PL APO 20X/0.80; HCX PL APO 40X/1.10 W CORR). Multispectral images were captured with microscope equipped with filter cubes for 6 dye separation and an external filter wheel (DFT51011). Image scanning was performed with 10% tiled image overlap. Z-stack imaging of 10 µm at 1.0 µm steps to cover the depth of the tissue.

### Data and code availability Padlock Probe Sequences

All data required to replicate the analyses, including mass spectrometry imaging data, H&E images, spaceranger output files, sequencing data, ISS data, Padlock Probe Sequences file, and additional files for all samples presented in this study will be available upon publication.

## Acknowledgements

The study was supported by Swedish Foundation for Strategic Research, Knut and Alice Wallenberg Foundation (KAW 2018.172) and Science for Life Laboratory. JL, MN was funded by grants from the Swedish research council (Dnr: 2022-03984, 2020-06182) and EU H2020 project EASI-Genomics. We would like to thank the National Genomics Infrastructure (NGI), Sweden for providing support, Drs Lukas Käll, Amelia Parker, and Xesús Abalo for input on the manuscript.

## Author contributions

MV, JL, PA initiated the project. MV, RM, AN, RS, HL performed the experiments. MV, LL, AN, PB, JF, PC analyzed the data and generated the figures. ME revised the statistical analysis. XZ, PS, provided model system and human samples. MV, RM, JL, drafted the manuscript with input from other authors; all authors read and approved the final manuscript. JL and PA provided project guidance and supervision.

## Conflict of interest

MV, RM, LL, and JL are scientific consultants for 10x Genomics Inc. The remaining authors declare no competing interests.

## Supplementary Figures

**Suppl. Fig. 1.**
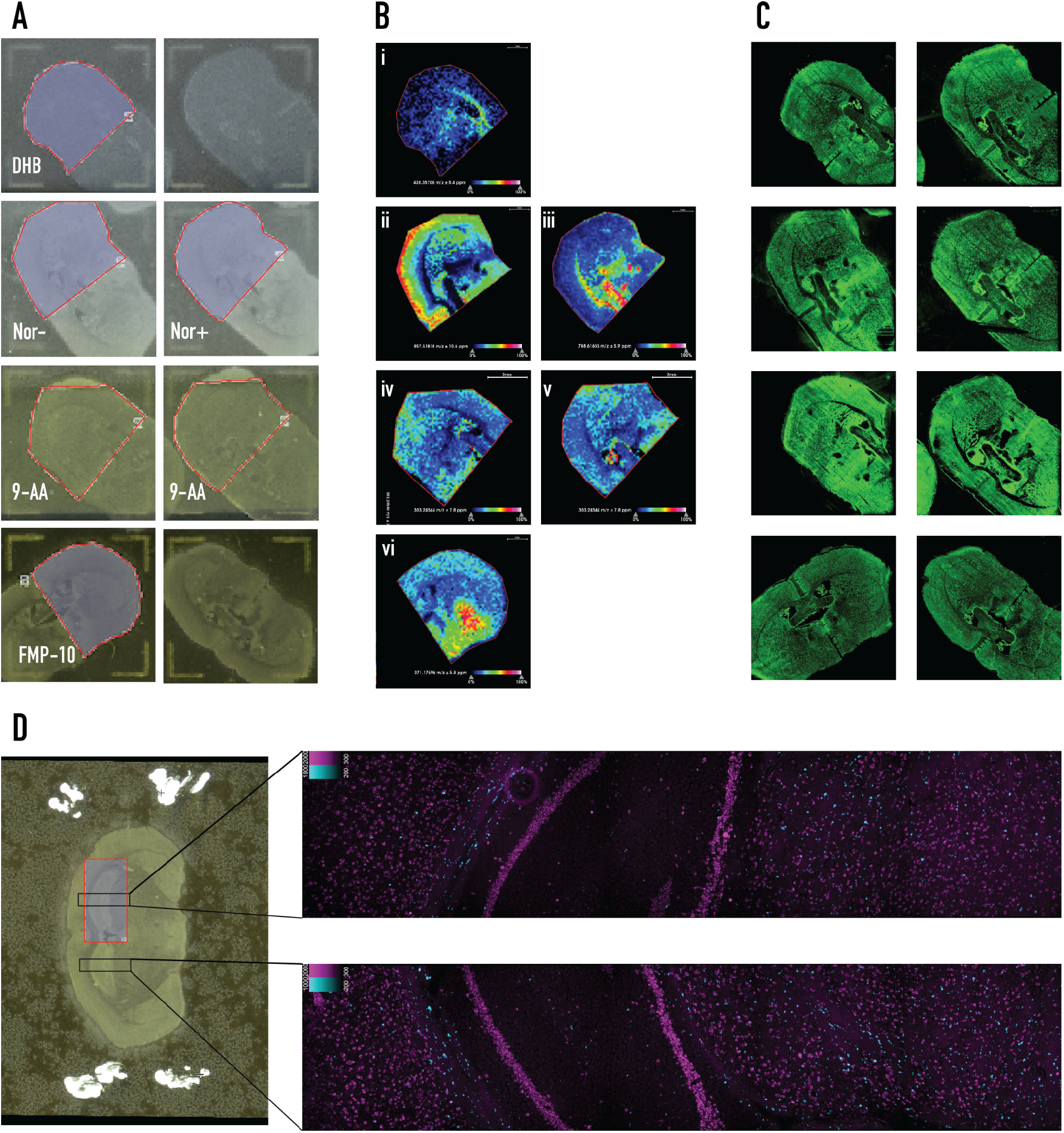
SMA using four different MALDI matrices. **A)** Mouse brain tissue sections from the striatal level were mounted onto a Visium Tissue Optimization slide and sprayed with four different MALDI matrices (DHB, Norharmane (analyzed in both positive and negative mode, shown as Nor+ and Nor-), 9-AA and FMP-10). Areas delimited by red lines: regions of interest imaged with MSI. **B)** Representative MSI results from: i) *m/z* 426.36, C18:1 L-Carnitine (DHB); ii) *m/z* 857.52, PI(36:4) (Nor-); iii) *m/z* 788.62 PC(36:1) (Nor+); iv,v) *m/z* 303.24, arachidonic acid (9-AA); vi) *m/z* 371.17, GABA (FMP-10). Nor+ and Nor-: Norharmane analyzed in positive and negative mode, respectively. Positive and negative modes produce positively and negatively charged ions and analyzed respectively. **C)** Fluorescence microscopy images of mRNA footprint captured with polydT probes after MALDI-MSI. **D)** Targeted In Situ Sequencing data demonstrate similar rolling circle product (RCP) density generated from MALDI-MSI processed region (upper right panel) and non-processed region (lower right panel) for demarcated regions of interest in the mouse coronal section. Targeted ISS simultaneously probed for housekeeping gene, *Gapdh* labelled in Magenta (Cy5), and a panel of five control genes - *Foxj1, Plp1, Lamp5, Rorb* and *Kcnip2* that are labelled in Cyan (AF750).

**Suppl. Fig. 2.**
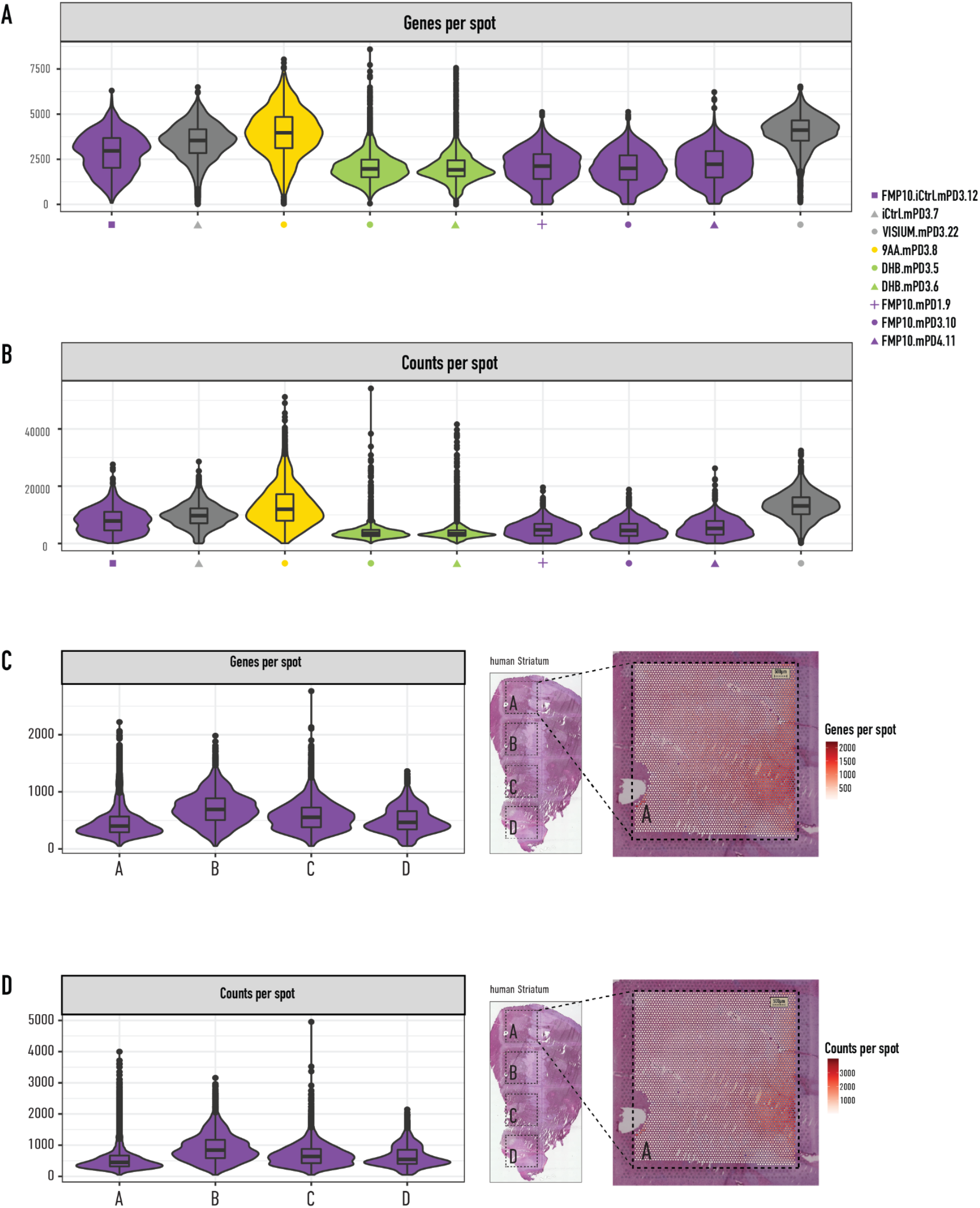
Spatial transcriptomics data quality control. **A)** Violin plots and box plots illustrating the number of unique genes per spot across technical and biological conditions of the mouse striatum data. **B)** Violin plots and box plots illustrating the number of unique molecular identifiers (UMIs) per spot across technical and biological conditions of the mouse striatum data. **C)** Violin plots and box plots illustrating the number of unique genes per spot of the human striatum data. The human sample H&E was used as a legend to indicate the four capture areas A-D. On the right, spatial featureplot representing the number of genes per spot of a representative capture area (i. e. capture area A). **B)** Violin plots and box plots illustrating the number of unique molecular identifiers (UMIs) per spot of the human Striatum data. On the right, spatial featureplot representing the number of UMIs per spot of a representative capture area (i. e. capture area A).

**Suppl. Fig. 3.**
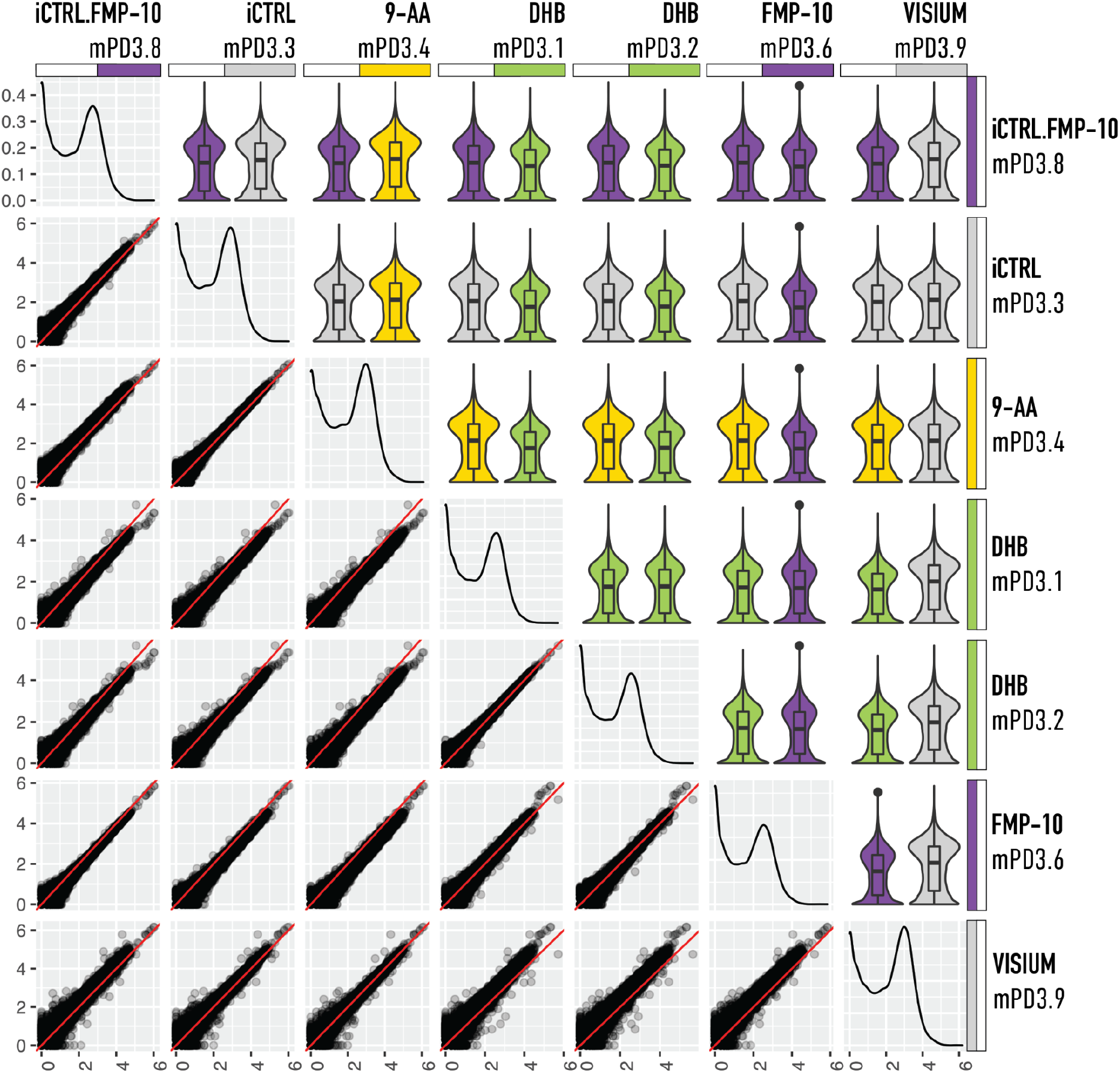
Pairwise scatterplots and violin plots of gene to gene log UMI count across technical conditions. Consecutive striatum sections of the same mouse (mPD3) were used in this analysis. The red line highlights a 1-to-1 relationship.

**Suppl. Fig. 4.**
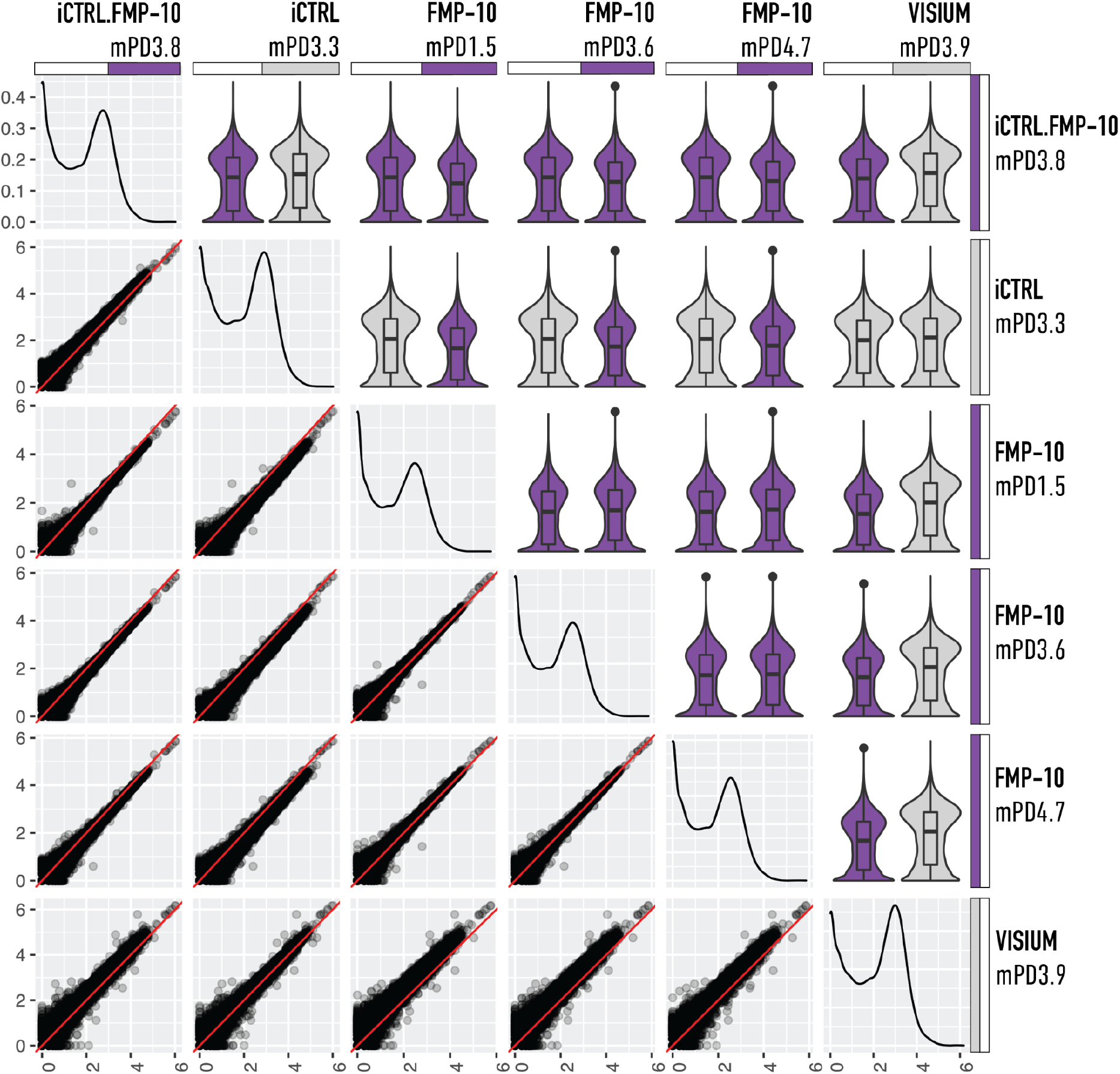
Pairwise scatterplots and violin plots of gene to gene log UMI counts across biological replicates. Several striatum sections of three different 6-OHDA mice (mPD1, mPD3, mPD4) were used in these analyses. The red line highlights a 1-to-1 relationship.

**Suppl. Fig. 5.**
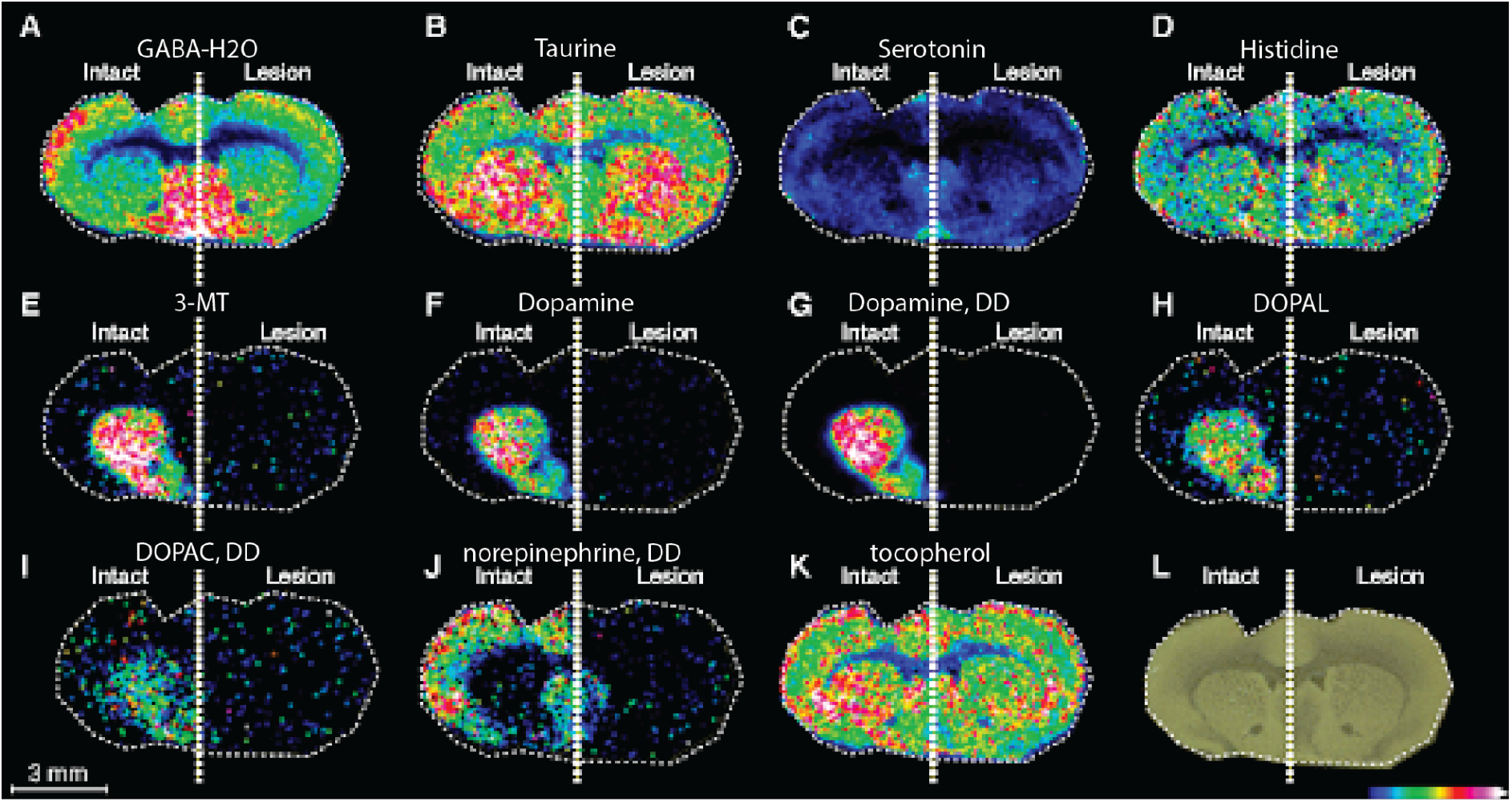
Selection of ion images from key neurotransmitters and metabolites in coronal tissue sections from a mouse PD model. Ion distribution images of (A) GABA-H2O, (B) Taurine, (C) Serotonin, (D) Histidine, (E) 3-MT, (F) Dopamine, (G) double derivatized (DD) Dopamine, (H) DOPAL, (I) double derivatized (DD) DOPAC, (J) double derivatized (DD) norepinephrine, (K) tocopherol, and (L) scanned image of the coronal mouse tissue section that was analyzed. All ion distribution images are scaled to 50% of max intensity and presented as single derivatized species, unless otherwise stated.

**Suppl Fig. 6.**
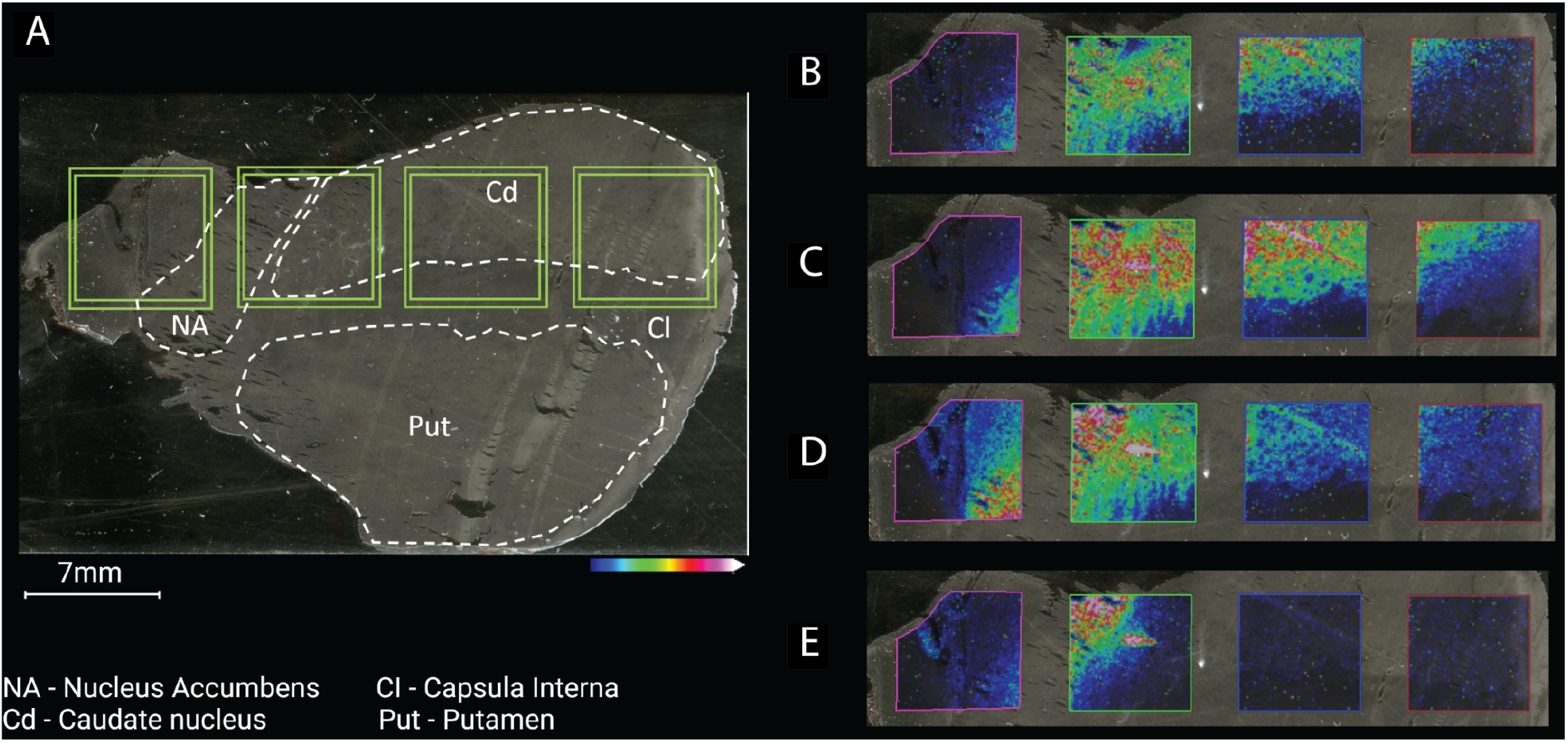
A scanned image of the human striatal tissue section on a Visium glass slide. (**A**) Whole tissue scan with annotated brain regions, where green squares indicate the areas coated with oligonucleotides. Ion images of **(B)** dopamine, **(C)** 3-MT, **(D)** serotonin, and **(E)** norepinephrine (double derivatized). All ion distributions are scaled to 50% of the maximum intensity and are all displayed as single derivatized species, unless otherwise stated.

